# Accessible spatial host–microbe profiling in tumour tissues

**DOI:** 10.64898/2026.07.02.735997

**Authors:** Yuk Kei Wan, Amanda Ng, Jing Yang Tham, Bo Li, Lim Gek Kiang Michelle, Iain Tan, Shyam Prabhakar, Niranjan Nagarajan, Grace Yeo, Minghao Chia

## Abstract

The ability of current spatial transcriptomics platforms to sensitively and specifically detect tissue-resident microbes alongside the whole host transcriptome remains limited. Here we present HOst MicrobE Spatial-seq (HOMES-seq), a Visium-based workflow for joint spatial profiling of microbial species and the host transcriptome in formalin-fixed paraffin-embedded (FFPE) colorectal tumours. HOMES-seq incorporates an analytical framework to distinguish contaminant-derived signals from bona fide tissue-resident microbes while improving detection sensitivity and reducing sequencing costs relative to existing approaches.

## Main text

Despite recent advances in highly multiplexed spatial detection of microbes, widespread adoption remains limited by requirements for specialized instrumentation and bespoke workflows. Imaging-based approaches, including par-seqFISH, HiPR-FISH and bacterial MERFISH, rely on complex setups such as microfluidics and expansion microscopy, and barcode constraints limit simultaneous profiling of microbes and the host transcriptome^1–3^. Spatial sequencing methods such as SHM-seq and SmT require custom slide chemistries bearing barcoded primers for microbial (16S/18S/ITS) and host (oligo-dT) capture that are not yet manufactured at scale^4,5^. There is therefore a need for accessible, out-of-the-box approaches to resolve spatial host–microbe interactions across tissues such as the gastrointestinal tract and skin. This need is particularly acute in colorectal cancer (CRC), where tumour-associated microbiomes can promote oncogenic signalling and tumour-promoting inflammation^6^. An ideal method would sensitively detect microbial species alongside the whole host transcriptome, achieve a high signal-to-noise ratio, be compatible with formalin-fixed paraffin-embedded (FFPE) tissues to leverage clinical archives, and avoid specialized equipment. To address this gap, we developed HOMES-seq, a spatial-omics workflow built using commercially available reagents and standard Visium infrastructure without modification of existing spatial chips that enables joint profiling of host transcripts and microbial species together with an analytical framework for contaminant identification.

To enable sensitive detection of tumour-associated microbes in human FFPE tissue sections, we developed a custom probe design pipeline targeting microbial genera and species enriched in CRC (Fig. 1a, Supplementary Table 1). A key challenge for spatial profiling of microbes is designing species-specific probes against highly conserved microbial ribosomal RNAs (rRNAs). To address this, we developed a computational pipeline for identifying microbial probe pairs compatible with the Visium FFPE workflow, targeting a panel of gut- and colorectal cancer (CRC)-associated genera and species (https://github.com/Chiamh/visium_probe_design_nf; Supplementary Table 1). The pipeline extracts 16S and 23S rRNA sequences from NCBI reference genomes and generates tiling 25-mer probes that are filtered based on GC content, melting temperature (Tm), and sequence similarity to curated SILVA databases (version 138.2) using BLASTN^7^. Because microbial rRNAs exhibit substantial sequence conservation, individual 25-mer probes frequently showed predicted off-target binding, defined as alignments with fewer than five mismatches. We therefore leveraged the paired-probe architecture of the Visium FFPE assay and inferred target specificity from the intersection of candidate matches for adjacent probe pairs, substantially reducing the likelihood of mis-hybridization (Fig. 1a). Where possible, tumour-associated species were supported by concordant signal across multiple independent probe pairs and by probes whose predicted off-targets correspond to phylogenetically distant or environmentally restricted taxa unlikely to be present in human tissue (Supplementary Table 1). Genus-level probes were additionally included for selected taxa when predicted hybridization patterns spanned multiple closely related species (Supplementary Table 1). Probe pairs were prioritized according to predicted specificity and thermodynamic properties (Methods) and integrated with the Visium Human Transcriptome Probe Kit v2.0 to enable joint spatial profiling of microbial species and host gene expression.

**Figure 1:**
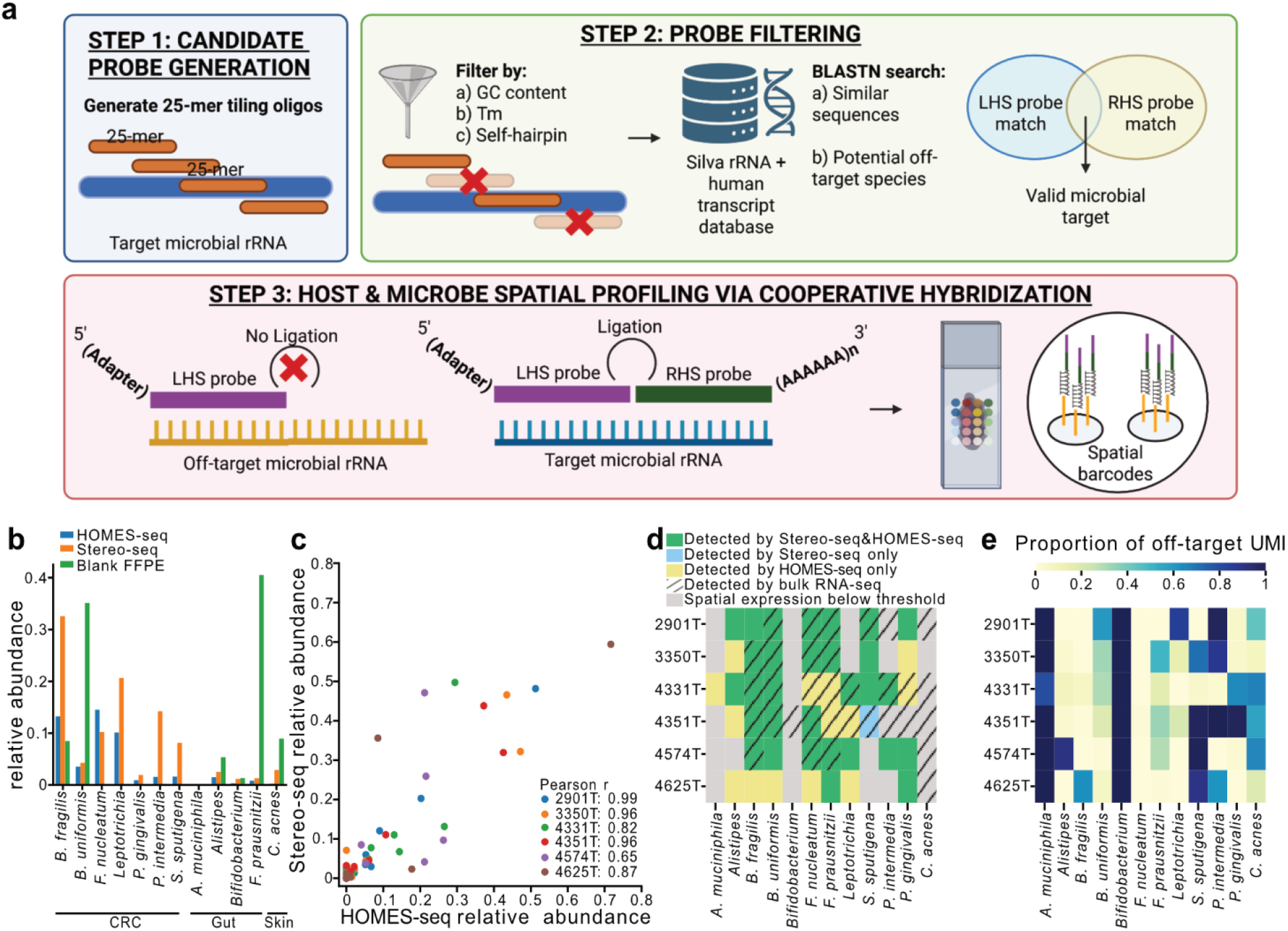
Probe-based HOMES-seq is concordant with the sequencing-based spatial platform Stereo-seq v1 for microbial transcript detection in FFPE sections. **a**. Schematic of the workflow for designing custom microbial probes and detection principles for HOMES-seq. 16S and 23S rRNA sequences from Colorectal Cancer -associated microbes are tiled into 25-mer candidate probe sequences. Probes are filtered based on GC content and desirable thermodynamic properties, then blasted against a database with microbial rRNAs and the human transcriptome to identify potential matches and off-targets. Due to high sequence similarity in rRNAs, individual probes often lack sufficient specificity. Species or genus specificity is inferred from the intersection of candidate matches for each probe pair, where two adjacent 25-mers must hybridize cooperatively for effective ligation and spatial barcoding, reducing the risk of artefactual species calls. **b**. Relative abundance of microbial species detected in tissue samples subjected to HOMES-seq (blue), Stereo-seq V1(orange), and blank FFPE samples subjected to bulk RNA-sequencing. **c**. Correlation of HOMES-seq and Stereo-seq V1 relative abundance. **d**. Microbial species detected by HOMES-seq and Stereo-seq V1 (green), Stereo-seq V1 only (blue), HOMES-seq only (yellow), or considered false positives due to low spatial autocorrelation similar to *C. acnes*, a contaminant (grey; Z test of Moran’s I, FDR < 0.01). Microbial presence in matched bulk RNA-sequencing is also shown (hatch). **e**. their proportion of off-target reads.

We next compared the performance of HOMES-seq to Stereo-seq-V1 (Stereo-seq OMNI V1.1)^8^, currently the only commercially available spatial sequencing platform enabling joint profiling of microbial species and the whole host transcriptome in FFPE tissues, in colorectal cancer tumours. Six tumour sections were serially sectioned and processed in parallel (Supplementary Fig. 1). When comparing signals from area-normalized spatial units (55 μm), HOMES-seq achieved substantially higher sequencing saturation despite having >10-fold lower sequencing requirements compared to Stereo-seq-V1 (10^8^ vs 10^9^ paired-end reads; Supplementary Table 2). At these sequencing depths, all HOMES-seq samples reached >80% saturation with ∼50,000 reads per spot. In contrast, while most Stereo-seq-V1 samples were also sequenced to near saturation, all had relatively few (5,000-25,000) UMI counts per spot due to most reads failing to map uniquely to the transcriptome (Supplementary Fig. 1a–b; Supplementary Table 2). Accordingly, HOMES-seq yielded a higher median number of detected genes per spot compared to Stereo-seq-V1 (8,000 vs 6,000). Notably, genes undetected by Stereo-seq-V1 were not limited to low-abundance transcripts. HOMES-seq consistently showed higher per-spot expression for canonical epithelial, fibroblast and neutrophil markers across a range of filtering thresholds (≥ 25, 40 or 50 non-zero spots), whereas Stereo-seq-V1 failed to detect some key host markers such as *COL3A1* in fibroblasts (Supplementary Fig. 1e; Supplementary Table 3).

In addition to host gene detection, we compared HOMES-seq and Stereo-seq-V1 for the detection of tumour-associated and gut microbial species. A major challenge in tumour microbiome studies is distinguishing bona fide tissue-resident microbes from environmental contaminants^9^. We reasoned that contaminant-derived signals would disperse during tissue permeabilization and library preparation, resulting in diffuse spatial distributions, whereas true tissue-resident microbes would exhibit localized enrichment. Consistent with this, the skin-associated bacterium *Cutibacterium acnes*, which is not expected to colonize CRC tumours, exhibited uniformly distributed signals with low spatial autocorrelation across tissue sections (mean Moran’s I = 0.12; Supplementary Fig. 3). In contrast, CRC-associated microbes displayed heterogeneous spatial autocorrelation across tumours, reflecting variable colonization patterns (Figure 1b). In tumours where these microbes were enriched, they formed spatially localized foci with significantly higher Moran’s I values than *C. acnes* (Z-test, FDR < 0.05), exemplified by *Leptotrichia trevisanii* in CRC4331T and CRC4574T (Supplementary Fig. 3). Because artefactual signals have also been reported to accumulate at tissue edges^10^, we implemented a two-step filtering framework that retained microbial signals with significant spatial autocorrelation followed by correction for tissue-edge artefacts using edge subtraction (Supplementary Fig. 4 and Methods).

To further assess background contamination, we sequenced tissue-blank FFPE samples processed in the same histopathology laboratory and compared their taxonomic profiles with spatial datasets following filtering (Fig. 1b). Several CRC-associated microbes including *Selenomonas sputigena, Prevotella intermedia, Porphyromonas gingivalis, Leptotrichia trevisanii* and *Fusobacterium nucleatum* were detected exclusively in tumour samples by both spatial platforms and were absent from blank controls (Fig. 1b). Although *Bacteroides fragilis* and *Bacteroides uniformis* were detected in both tumour and blank samples, their relative abundances differed significantly (p-value 3.46E-3) from those of common background taxa, including the reported beneficial commensal gut microbes *Faecalibacterium prausnitzii, Bifidobacterium spp., Alistipes spp*. and the skin microbe *C. acnes*. Furthermore, the marked difference in spatial distribution between CRC-associated microbes and the common background taxa (p-value HOMES-seq: 6.24E-3, Stereo-seqV1: 8.75E-2) supports the CRC-associated microbes’ classification as tumour-associated signals rather than laboratory contaminants (Fig. 1b; Supplementary Fig. 3). Together, these analyses demonstrate that integrating negative controls with spatial patterning provides an effective strategy for distinguishing biological microbial signals from contaminant-derived artefacts in CRC tissues.

Off-target hybridization can introduce chimeric, false-positive signals in probe-based spatial transcriptomics platforms^11^. We therefore evaluated the extent of such events in HOMES-seq by quantifying the proportion of reads classified as chimeric or partially mapped. Across all microbial targets, these events were strongly enriched among low abundance detections (Figure 1c and Supplementary Figure 1f). Taxa supported by fewer than 100 UMIs consisted predominantly of chimeric reads (median, 100%), consistent with rare stochastic probe mis-hybridization events. In contrast, microbial targets with greater read support exhibited consistently low chimeric rates (median, 1.31% among species supported by >500 UMIs) (Supplementary Figure 1f). These findings indicate that HOMIES-seq exhibits minimal off-target hybridization, with artefactual signals largely restricted to low-support detections and readily distinguishable from high-confidence microbial signals.

We next evaluated whether the resulting microbial profiles were concordant across HOMES-seq, Stereo-seq-V1 and bulk metatranscriptomes from matched tissue blocks. Relative abundances of CRC-associated microbes were highly concordant between HOMES-seq and Stereo-seq-V1 in all six samples (Pearson’s R 0.61 - 0.99; p-value 2.91E-8 - 4.5E-2; Fig. 1d). The weakest correlation was observed in CRC4574T due to disproportionately high *Leptotrichia* abundance detected by Stereo-seq-V1. Relatively abundant CRC-associated microbes, including *Leptotrichia trevisanii, Fusobacterium nucleatum, Bacteroides fragilis* and *Bacteroides uniformis*, were consistently detected across all three assays (Spearman ρ: 0.29, p-value: 6E-4, Fig. 1e). Occasional discrepancies between spatial technologies and bulk metatranscriptomics were attributable to technical limitations. For example, in CRC3350, *B. fragilis* and *F. nucleatum* were detected by both spatial platforms but not by bulk metatranscriptomics because of low RNA yield (Fig. 1e and Supplementary Table 2). Detection of lower-abundance taxa (% by bulk) such as *Alistipes spp*. and *Porphyromonas gingivalis* was less consistent between bulk and spatial assays, likely reflecting stochastic sampling effects. The spatial distribution of microbial-positive regions detected by both HOMES-seq and Stereo-seq-V1 broadly agreed with orthogonal RNAscope staining patterns (Supplementary Fig. 2). However, HOMES-seq detected microbial neighbourhoods in sparser regions, consistent with improved sensitivity despite substantially lower sequencing requirements (10^8^ versus 10^9^ paired-end reads; Supplementary Table 1). Together, these findings indicate that HOMES-seq enables more sensitive and spatially resolved microbial detection while maintaining high-quality host transcriptome profiling at substantially reduced sequencing cost compared to Stereo-seq-V1.

Transcriptome-wide host profiling by HOMES-seq enables spatial comparisons of microbial neighbourhoods with local immune and stromal signalling programmes. Previous studies have shown that tumour niches that are colonized by CRC-associated microbes were also enriched in myeloid cells such as tumour-associated neutrophils (TANs)^12–14^ . To investigate this, we quantified the spatial enrichment of host immune programmes relative to defined microbial neighbourhoods within each tumour section. Agreeing with the reported chronic inflammation feedback loop among IL1B, inflammatory cancer associated fibroblast (iCAFs), and neutrophil activation^15^, we-observed that regions expressing *IL-1B* and neutrophil-recruitment factors (*CXCL8, CXCL1, ATF3, TIAM2*), markers for iCAFs (*MMP1, MMP3*) and the matrix metalloproteinase 9 (*MMP9*) were spatially adjacent in 5 of the 6 tumours (excluding CRC4331T, Moran’s I 0.408 - 0.701; p-value 0.001) (Supplementary Figure 5). This spatial coupling between microbial signals and immune activation is consistent with biologically structured host–microbe organization rather than background contamination.

The transcriptome-wide host coverage of HOMES-seq also facilitates systematic interrogation of immune functional states near microbe-associated regions for hypothesis generation and discovery. In particular, the proximity of TANs to microbe-rich regions raises the question of how neutrophil heterogeneity contributes to these spatial niches^16,17^. Through HOMES-seq, we quantified expression of a panel of genes reported to be associated with neutrophil extracellular trap (NET) formation, a neutrophil activation programme implicated in sustaining inflammatory tumour microenvironments and promoting oncogenic signalling^18–22^. We observed enrichment of a NET-associated transcriptional programme (*FPR1, S100A12, PDE4B, CSF3R, VNN3* and *PADI4*) in 2 of the 6 tumours (CRC2901T and CRC3350T; p-values 0.001) showing the strongest localized IL1B signalling proximal to microbial regions (Supplementary Fig. 6). To resolve neutrophil populations associated with bacterial abundance at single-cell resolution, we further complemented the spatial transcriptome data generated by HOMES-seq with Xenium profiling of matched tumour sections using a custom panel targeting ∼5,000 human genes together with 16S/23S rRNA sequences from 22 bacterial species and genera (Methods). Xenium-derived markers for TAN subtypes were further confirmed in both our internal single-cell RNA-seq data^23^ as well as a recently published pan-cancer neutrophil atlas^24^. We identified two transcriptionally and spatially distinct neutrophil populations: a *S100A8+/S100A9+* population representing an immature or primed inflammatory state and a *HCAR2+/IL-1B+* population with an activated phenotype proximal to microbial regions in 4 of the 6 tumours (Figure 2a-b). Scoring of various gene signatures in HOMES-seq data with bivariate Moran’s I analysis showed the same spatial patterning, with bacteria most proximal to activated TANs, which were in turn proximal to *MMP1+/MMP3+* iCAFs, suggesting a pro-tumour inflammatory axis accompanied by matrix remodelling (Figure 2c, Supplementary Figure 5, Supplementary Table 4). HOMES-seq’s revealing the co-localisation of microbes and the IL-1B-iCAF-neutrophil-created immunosuppressive environment sheds light to potential targets to improve immunotherapy efficacy. These results highlight the utility of HOMES-seq for functional inference of immune states from spatial transcriptomes, and discovering spatially coordinated immune activation states in proximity to CRC-associated microbial neighbourhoods *in situ*.

**Figure 2:**
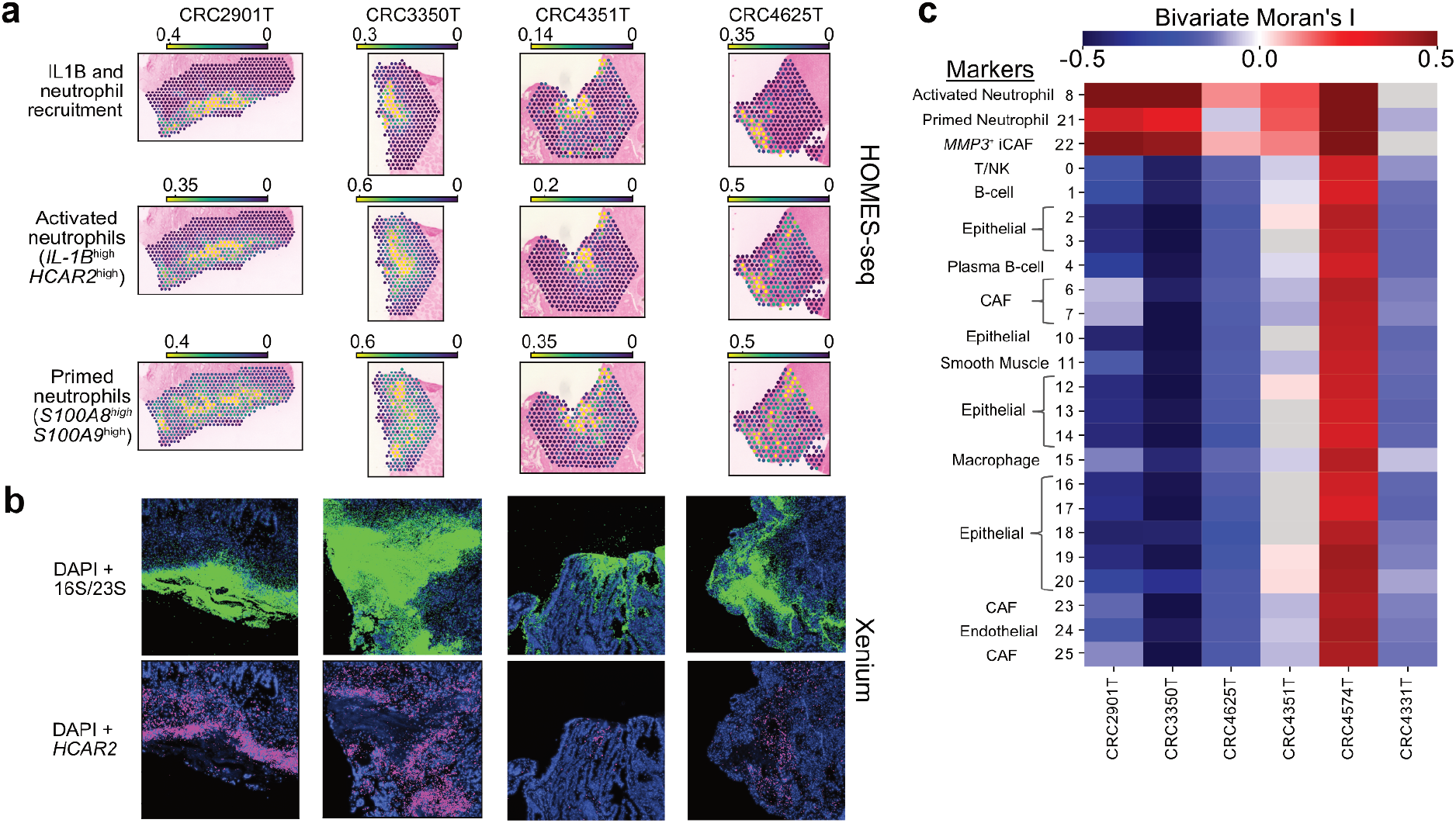
HOMES-seq identifies spatially distinct primed and activated neutrophil populations near microbial colonisation in CRC tumors. **a**. HOMES-seq-based gene set scores for IL-1B and neutrophil recruitment (top) and distinct neutrophil subtypes. **b**. Xenium-based DAPI and microbial 16S/23S rRNA signals (top) and DAPI and HCAR2 signals (bottom). c. Heatmap showing the Bivariate Moran’s I spatial auto-correlation between microbes and various cell-type specific marker transcripts.

We present HOMES-seq, a Visium-based workflow that enables joint profiling of host gene expression and microbial species in FFPE tissues. HOMES-seq is compatible with archived clinical samples, provides transcriptome-wide host coverage, and does not require specialized instrumentation or extensive modification of existing spatial transcriptomics capture arrays, thus addressing key limitations of existing methods. In comparison with Stereo-seq-V1, the only other commercial alternative for joint host-microbe profiling at present, HOMES-seq achieved improved sequencing efficiency and sensitivity, with higher gene and microbial detection at substantially lower sequencing depth. This cost-effectiveness is supported by the use of customizable probe sets for targeted microbial detection, which can be flexibly designed and synthesized at low cost. Low rates of off-target hybridization, assessed through read chimerism, together with concordance across orthogonal methods and the use of negative FFPE controls and spatial patterning, support the specificity of detected microbial signals. Leveraging whole-transcriptome information, we identified significant spatial enrichment of CRC-associated microbes and neutrophil-enriched host neighbourhoods, accompanied by iCAFs, activated *HCAR2+/IL-1B+* TANs and matrix-remodelling functions. These findings highlight the ability of HOMES-seq to resolve spatial host–microbe interactions within their native tissue context. Notably, the HOMES-seq workflow is directly compatible with higher-resolution platforms such as Visium HD, allowing finer-scale characterization of host–microbe interactions. Together, HOMES-seq provides a scalable and accessible approach for interrogating spatial host–microbe interactions in tumours, expanding the toolkit for studying cancer microbiomes.

## Methods

### Participant recruitment

Participants were recruited by the National Cancer Centre Singapore (NCCS) with Institutional Review Board approval (IRB reference no. 2018-2795). All associated protocols for this study conducted at the Genome Institute of Singapore (GIS) were approved by the Agency for Science, Technology and Research Institutional Review Board (A*STAR IRB reference no. 2024-016).

### Formalin fixed & paraffin embedded (FFPE) tissue preparation

Colorectal cancer tissues were fixed in formalin and embedded in paraffin according to the “Visium CytAssist Tissue Preparation Guide” (CG000518 Rev C).

### RNA Quality Assessment (DV200)

RNA was extracted from 10 μm tissue rolls using the RNeasy FFPE Kit (Qiagen, Cat. No. 73504), according to the manufacturer’s protocol. To assess the RNA quality, RNA ScreenTape (Agilent Technologies, Cat. No. 5067-5576) was used to determine DV200 values (>30%).

### Tissue sectioning for Spatial Assays

Tissue blocks were sectioned at 5 μm thickness and placed on the recommended slides for each assay. Serial tissue sections were taken for 10x Xenium, 10x Visium, Stereoseq and H&E staining, where possible. Tissue slides were dried according to the user guides of each assay..

### Screening of bacterial signal using Single molecule Fluorescence In Situ Hybridisation (smFISH) assay

Tissue slides were baked at 60 °C for 1h and cooled to room temperature for 5 min. Deparaffinisation was carried out by submerging slides in Xylene jar 1 for 5 min, Xylene jar 2 for 5 min, 100% Ethanol jar 1 for 5 min, 100% Ethanol jar 2 for 5 min, 95% Ethanol jar 1 for 5 min, 95% Ethanol jar 2 for 5 min, 70% Ethanol for 5 min. Slides were blotted dry in between each step, and were stored in 2X Saline-sodium citrate (SSC) buffer at room temperature before probe hybridisation.

Universal bacterial probes (EUB338)^25^ and probes specific for *Fusobacterium nucleatum* (FUS714)^26^ were ordered from Integrated DNA Technologies with a fluorophore attached at the 5’ end (Supplementary Table 5). Probes were diluted to 2 μM in warm (47°C) hybridisation buffer (20mM Tris-HCl pH 8.0, 0.9M NaCl, 0.01% SDS, 0.2 μm filtered 10% Formamide) Hybridisation was carried out in an opaque humidified pipette tip box, lined with a piece of parafilm and 100 μl of pre-warmed probe mix was pipetted onto the parafilm. Tissue slides were gently placed over the probe mix (tissue face down) and were incubated for 1.5-2 h at 47 °C in the dark.

Slides were carefully removed and washed three times with FISH wash buffer (225mM NaCl, 20mM Tris-HCl pH 8.0, 5 mM EDTA, 0.2 μm filtered) at 48 °C, for 5 minutes per wash. Slides were blotted dry and mounted with ProLong™ Gold Antifade Mountant with DNA Stain DAPI (Invitrogen™, Cat. No. P36935) and sealed with coverslips. Slides were cured overnight in the dark at room temperature.

### Chromogenic detection of bacterial signal using RNAScope

RNAScope assay was performed with RNAScope®2.5 HD Duplex Detection Kit (Chromogenic) (Advanced Cell Diagnostics, Inc., Cat. No. 322430 and 322500) with RNAscope Probe-EB-16S-rRNA, C1 (Advanced Cell Diagnostics, Inc., Cat. No. 464461) and RNAscope Probe - B-Fusobacterium-23S-3zz-C2 (Advanced Cell Diagnostics, Inc., Cat. No. 486411-C2). Tissue slides were pretreated according to the user manual “Formalin-Fixed Paraffin-Embedded (FFPE) Sample Preparation and pretreatment” (ACDBio Document No. 322452), with slight modifications to the set up for target retrieval by using a heat-resistant plastic rack and water bath in place of a steamer.

### 10x Visium FFPE

Tissue slides were prepared according to “Visium CytAssist for FFPE – Deparaffinization, H&E Staining, Imaging & Decrosslinking (CG000520 Rev B)” and “Visium CytAssist Spatial Gene Expression Reagent Kits User guide (CG000495 Rev E)”. Custom probes were ordered, prepared and added into the assay according to “Technical Note for Custom Probe Design (CG000621 Rev D).

### Haematoxylin and Eosin staining and Imaging

Tissue sections were stained with haematoxylin and eosin (H&E) using a Leica HistoCore SPECTRA ST automated stainer (Leica Biosystems, Wetzlar, Germany) following manufacturer’s instructions. Briefly, sections underwent sequential deparaffinisation in xylene, rehydration through a graded ethanol series (100%, 95%, 70%), Harris haematoxylin staining with acid-alcohol differentiation and lithium carbonate bluing, followed by eosin counterstaining and dehydration through ascending ethanols and xylene prior to coverslipping. Stained slides were digitised on a Philips SG300 whole-slide scanner (Philips Digital Pathology, Amsterdam, Netherlands) at 0.25 μm/pixel (40× equivalent magnification) with continuous autofocus enabled, generating iSyntax files managed and reviewed within the Philips IntelliSite Pathology Solution.

### Custom microbial probe design for Visium FFPE

Probe pairs for each target microbial sequence comprised left-hand side (LHS) and right-hand side (RHS) half-probes of 25 nucleotides each, following design parameters adapted from the 10X Genomics Custom Probe Design Technical Note (CG000621), and implemented in a custom Nextflow pipeline (https://github.com/Chiamh/visium_probe_design_nf/). Candidate probe regions were generated using OligoMiner (version 1.7)^27^, which tiles all eligible 25-nucleotide windows across each target sequence subject to a GC content of 44–72% and a melting temperature (Tm) of 48–77 °C, calculated under Visium hybridisation conditions (1× SSC, 0% formamide, 165 mM Na^+^). Reference sequences were obtained from NCBI, where ribosomal RNA (rRNA) targets comprised 16S and 23S sequences for bacteria and 18S and 28S sequences for fungi. To assess off-target binding, candidate probes were aligned against a reference database comprising the human transcriptome, transfer RNAs, and small and large subunit rRNA sequences from the SILVA 138.2 database using BLASTN (version 2.12.0). Probes were retained only if off-target matches carried at least five mismatches in at least one of the two half-probes; where complementarity to an off-target exceeded 75% (>20 bp) in one half-probe, the pair was retained provided the partner half-probe showed no cross-hybridisation to the same off-target sequence. Thermodynamic filtering further required that self-hairpin and self-dimer Tm values did not exceed 45 °C and that the difference between the on-target hybridisation Tm and the self-hairpin Tm exceeded 10 °C to minimise self-inhibition. Probe pairs were required to be non-overlapping to prevent competition for adjacent binding sites. Candidates were first prioritised for species- or genus-level specificity, then filtered on the thermodynamic criteria above, and final probe sequences were appended with Visium adapter sequences for downstream spatial transcriptomics workflows (Supplementary Table 1).

### Xenium sample preparation and imaging

Xenium prime assays were performed according to 10X Genomic User Guides (CG000578, CG000760), using the pre-designed 5000-gene panel and an additional 100 custom genes. After the Xenium run was completed, sample sections were retrieved for downstream H&E. The following 10X Genomics reagents and accessories were used to perform the assays: Xenium Prime 5K Human Pan Tissue and Pathway Assay Kit (1000671), Xenium Prime 5K Add-on Custom 51-100 Gene Panel (1000766), Xenium Thermocycler Adaptor v2 (1000739).

In addition to 67 human genes, 33 custom probes were designed targeting the 16S and 23S regions of 22 bacterial species (Supplementary Table 6). Probes were filtered against an off-target database consisting of the human transcriptome and SSU and LSU sequences from SILVA (v 138.1, NR99 trunc). Detailed Xenium probe filtering and thermodynamic selection parameters are proprietary to 10x Genomics and are not publicly available.

### Spot-level filtering for Stereo-seq and Visium, and Xenium samples

Stereo-seq data was processed using the Stereo-seq Analysis Workflow (SAW v8.1.0) which removed all out of tissue spots. Visium data was processed using Space Ranger (v 3.1.3), and spots within tissue were retained by filtering away those with fewer than 300 UMI counts.

Raw Xenium data was generated using the Xenium Onboard Analysis pipeline (XOA v3.0.0.15). In the initial QC, cells with at least 5 genes and >10 total host transcriptome counts were retained. This resulted in a total of 1,074,275 cells retained across 4 samples, with a median of 241 counts and 191 genes detected per cell.

### Accessing assay quality and sensitivity of Stereo-seq and Visium

We compared the transcript detection sensitivity of Stereo-seq and Visium using three metrics, namely n-counts per gene, sequencing saturation, and major cell type gene marker expression. To match the spot size of Visium, we used the stereopy package to simulate bin-220 (approximately 55μm diameter spots) for Stereo-seq samples for the following comparisons. The following subsections details how we obtain quality metrics from the samples:

#### 1) Sequencing quality metrics and sequencing saturation

For each sample, we obtained sequencing quality metrics, including total counts and n-counts per gene, with scanpy v1.11.4’s^28^ pp.calculate_qc_metrics function. To calculate sequencing saturation for each sample, we randomly selected a set number of reads per spot and calculated sequencing saturation per spot with the formula: sequencing saturation = 1-(unique genes in the spot/total number of reads in the spot). We visualised the sequencing saturation with scatter plots using the matplotlib^29^ package and a trendline using the make_interp_spline function from the scipy package.

We compared the n-count per gene, which is an assay sensitivity estimate, across the samples, using seaborn v0.13.2’s^30^ distplot function for visualization. We compared the n-count per gene for the whole transcriptome for Stereo-seq and Visium. We visualised the n_count per gene with histograms plotted with the seaborn v0.13.2^30^ package and a vertical line indicating the median n-count per gene with the matplotlib^29^ package.

#### 2) Major cell type marker expression

Across the Stereo-seq and Visium samples, we compared the expression of gene markers from major cell types, namely epithelial cells (*EPCAM, CDH1, MUC2*), endothelial cells (*COL11A1, COL3A1, AEBP1*), fibroblast (*PLVAP, PECAM1, STC1*), and neutrophils (*ITGAX, CSF3R*). Using scanpy v1.11.4’s^28^ pp.log1p function, we log+1 transformed the gene expressions for each spot. For each sample, we kept spots with an expression larger than 0 for each gene, averaged the filtered spots’ expression for each gene, and visualised the log(average_spot_expression) with a box plot for each gene marker, if the gene marker has more than 50 non-zero spots, using seaborn v0.13.2’s^30^ boxplot and stripplot functions.

### Identifying microbial contaminant signals by edge substraction

For each spatial sample, we calculated the averaged microbe or species-specific expression on the tissue edge spots to account for potential microbial contaminations. Microbial species/signatures were subsequently decontaminated by subtracting, on a per-spot basis, the mean tissue-edge signal of the corresponding microbe.

### Evaluating off-target effects of Visium probes for microbes

For each Visium sample, 10x Genomics’ CellRanger software provides the mapping quality (mapq) in the alignment possorted_genome_bam.bam file. The four mapq values are 255 (both read halves map to the same probe), 3 (each read half maps to a different probe), 1 (one read half maps to a probe and the other half does not), and 0 (neither read half maps to a probe). The percentage of reads with a mapq of 3 indicates the formation of left-hand side (LHS) and right-hand side (RHS) probe chimeric, providing an estimate of off-target effects. Across the Visium samples, we calculated the proportion of off-target reads by dividing the sum of reads with mapq 3 and 1 by the total number of reads for each microbe detected. We then visualised the proportion of off-target reads and the log(total reads) with a scatter plot using the matplotlib^29^ package. We also visualised the proportion of off-target reads across microbes in each sample with a heatmap plotted using the seaborn v0.13.2^30^ package.

### Detecting microbial relative abundance in spatial samples

For each spatial sample, we calculated tissue-wide relative abundance for each microbial species detected in HOMES-seq. We first log10+1-transformed the HOMES-seq and Stereo-seq samples’ expression with scanpy v1.11.4’s^28^ pp.log1p function and calculated the mean log10 expression for each microbe. We decontaminated potential microbial contaminations as stated above. For each sample, we calculated the species-specific relative abundance by dividing the decontaminated and transformed species-specific expression with the sum of all microbe expressions, where the sum of all microbial relative abundance should equal 1. We also visualised the tissue’s local, bin-based expression for all microbes, *Fusobacterium nucleatum*, and *Bacteroides fragilis* with scanpy v1.11.4’s^28^ pl.spatial function.

### Processing of bulk RNA sequencing samples

We subjected rRNA-depleted RNA extracted from the six FFPE tissue blocks and blank FFPE blocks to bulk RNA sequencing. Briefly, we processed the samples by removing adapter sequences with fastp (version 0.22.0)^31^, aligning the fastq reads to the human genome (hg38) using STAR (version 2.7.9a)^32^, deduplicating reads with bbmap (version 38.93) clumpify.sh^33^, and performing taxonomic classification of unmapped, non-human reads with Kraken2 (version 2.1.2)^34^ using a 50 gb database built from RefSeq bacterial, archaeal, viral, fungal and human (hg38) genomes, as well as plasmid sequences.

### Evaluating potential microbial contaminations in spatial samples

To evaluate whether colorectal cancer and gut-specific species detected in HOMES-seq samples are contaminations, we subjected blank FFPE blocks to bulk RNA-sequencing. We processed the sequenced samples as described in the “Processing of bulk RNA sequencing samples” section. We calculated relative abundance for each species as follows, we normalised the Kraken-outputted abundance with the sum of Kraken-outputted abundances across all species detected using the customised Visium probes, where the relative abundances across all species detected by Visium probes sum up to 1. For each species detected by HOMES-seq, we compared and visualised the species’ relative abundance in HOMES-seq and Stereo-seq V1 samples to that in blank FFPE blocks with a bar plot using the matplotlib^29^ package.

We hypothesise that potential microbial contaminations would be spatially dispersed while microbes residing in the tumour would spatially cluster together. Hence, we calculated Moran’s I to access each microbial species spatial autocorrelation in both HOMES-seq and Stereo-seq V1 using the *squidpy v1.6.5*^35^ python package. For each species detected by HOMES-seq, we compared and visualised the species’ Moran’s I in HOMES-seq and Stereo-seq V1 samples with a bar plot using the seaborn v0.13.2^30^ package.

### Evaluating microbial abundance of spatial samples and bulk RNA sequencing

We evaluated whether the microbes called from samples subjected to spatial assays, 10X HOMES-seq or Stereo-seq V1, are also called using bulk RNA sequencing. For each sample, we collected total RNA from tissue shavings and depleted rRNA prior to subjecting the RNA sample to bulk RNA sequencing. We processed the sequenced samples as described in the “Processing of bulk RNA sequencing samples” section. For the bulk samples, we calculated microbe-only abundance by normalising reads assigned to each species with only non-human and non-unclassified reads; then, we defined expressed microbes as those whose microbe-only abundance has a non-zero uniform count ≥ 0.1. For spatial samples, we defined expressed microbes based on a one-tail z-test of the microbe’s Moran’s I and C. acnes Moran’s I, where confidently called microbes have a Benjamini-Hochberg adjusted p-value less than 0.01. Then, we visualised the microbes called among bulk, HOMES-seq, and Stereo-seq V1 using a heatmap plotted using the seaborn v0.13.2^30^ package.

### Distance-based gene set variation analysis in HOMES-seq samples

For each HOMES-seq sample, we first summed up all microbial expressions in each spot to obtain the per-spot expression for all microbes and determined spots with at least a 95 percentile expression of all microbes to define microbe-rich regions per section. We further utilised a custom script to remove individual highly expressed microbe-rich spots isolated from any clusters of highly expressed microbe spots. We used these defined microbe-rich areas as the starting area of the distance-based gene set variation analysis, where all the spots in these defined areas have a distance of 0.

We performed distance-based gene set variation analysis separately for microbe-rich regions, where we labelled the surrounding spots based on the number of spot(s) with range 1 to 10 spots from the defined regions. We use *FCGR3B, CSF3R* as canonical neutrophil gene markers and *EPCAM, CDH1, MUC2, and KRT20* for canonical epithelial cell gene markers. Investigating IL-1B-associated neutrophil recruitment, we calculated each spot’s mean expression of the gene set containing *IL-1B, CXCL8, CXCL1, ATF3*, and *TIAM2*. To normalise IL-1B-associated neutrophil recruitment with neutrophil and epithelial cell signatures, we divided each spot’s mean expression based on the IL-1B-associated neutrophil recruitment gene set with the spot’s mean expression based on canonical neutrophil and epithelial gene markers. We also calculated the normalised gene set mean expression of inflammatory cancer-associated fibroblast (iCAF) markers genes, containing FAP, ADAM12, GAS1, WNT5A, WNT2, IL7R, MMP1, CXCL12, and IGFBP5, and MMP9 gene expression by dividing the spot’s mean expression of fibroblast markers, including *COL11A1, AEBP1*, and *COL3A1*. We visualised the gene set mean expressions of IL1B-associated neutrophil recruitment, iCAF, and gene expression of MMP9 with scanpy v1.11.4’s^28^ pl.spatial function.

### Identifying spatially distinct neutrophil populations in Xenium samples

To annotate cell types within the Xenium data, we implemented a clustering pipeline using Scanpy^28^ (v1.11.5) and BANKSY^36^ (v1.3.4). Briefly, count data was log-normalized to a library size of 150. Then, spatial features were introduced using BANKSY^36^ (M = 0, k_geom = 15, lambda = 0.1), followed by dimensionality reduction via PCA. Batch correction was performed via Harmony^37^ (v0.2.0) using sample as the batch variable. A KNN graph (k = 20, PCs = 10) was built using the batch-corrected embeddings and Leiden clustering^38^ (resolution = 0.6) was applied to yield 26 clusters. Marker genes for each cluster were identified via the Wilcoxon rank-sum test. To facilitate annotation of major cell types within the spatial clusters, we compared the marker gene expression to our previously published scRNA-seq CRC atlas^23^. To further characterize the two neutrophil populations observed in our spatial data, we calculated correlations between the expression profiles of the neutrophil clusters from our spatial data, with the neutrophil sub-clusters from a reanalysis of the neutrophil population from our CRC atlas, as well as a public neutrophil scRNA-seq atlas^24^.

### Identifying spatially distinct neutrophil populations in HOMES-seq

For each tumour section processed with HOMES-seq, we calculated the mean gene set expression for the top 20 markers from each cluster defined from the Xenium samples and the mean microbial expression of the CRC associated microbes *B. fragilis, F. nucleatum*, and *Lepotrichia spp*. We then measured the correlation of the locations of microbes and cells from each Xenium-defined cluster with bivariate Moran’s I using the esda v2.9.0, pysal v4.14.1^39^, and squidpy v1.6.5^35^ packages. We visualised the bivariate Moran’s I across samples with a heatmap plotted with the seaborn v0.13.2^30^ package. We visualised the gene set mean expressions of neutrophil clusters 8 and 21 with scanpy v1.11.4^28^ pl.spatial function. Cluster markers used for this analysis are listed in Supplementary Table 4.

## Data availability

All HOMES-seq and Stereo-seq V1 samples will be available upon manuscript acceptance.

## Code availability

Source code for scripts used to analyze the data is available at https://github.com/CSB5/HOMES-seq. Code for the Probe-design Nextflow pipeline is available at https://github.com/Chiamh/visium_probe_design_nf.

